# DeeReCT-APA: Prediction of Alternative Polyadenylation Site Usage Through Deep Learning

**DOI:** 10.1101/2020.03.26.009373

**Authors:** Zhongxiao Li, Yisheng Li, Bin Zhang, Yu Li, Yongkang Long, Juexiao Zhou, Xudong Zou, Min Zhang, Yuhui Hu, Wei Chen, Xin Gao

**Affiliations:** King Abdullah University of Science and Technology (KAUST), Computational Bioscience Research Center (CBRC), Computer, Electrical and Mathematical Sciences and Engineering (CEMSE) Division, Thuwal, 23955-6900, Saudi Arabia; Department of Biology, Southern University of Science and Technology (SUSTech), Shenzhen, 518055, China; Cancer Science Institute of Singapore, Singapore 117599, Singapore

**Keywords:** Polyadenylation, Gene regulation, Deep learning, Bioinformatics

## Abstract

Alternative polyadenylation (APA) is a crucial step in post-transcriptional regulation. Previous bioinformatic works have mainly focused on the recognition of polyadenylation sites (PAS) in a given genomic sequence, which is a binary classification problem. Recently, computational methods for predicting the usage level of alternative PAS in a same gene have been proposed. However, all of them cast the problem as a non-quantitative pairwise comparison task and do not take the competition among multiple PAS into account. To address this, here we propose a deep learning architecture, DeeReCT-APA, to quantitatively predict the usage of all alternative PAS of a given gene. To accommodate different genes with potentially different numbers of PAS, DeeReCT-APA treats the problem as a regression task with a variable-length target. Based on a CNN-LSTM architecture, DeeReCT-APA extracts sequence features with CNN layers, uses bidirectional LSTM to explicitly model the interactions among competing PAS, and outputs percentage scores representing the usage levels of all PAS of a gene. In addition to the fact that only our method can predict quantitatively the usage of all the PAS within a gene, we show that our method consistently outperforms other existing methods on three different tasks for which they are trained: pairwise comparison task, highest usage prediction task and ranking task. Finally, we demonstrate that our method can be used to predict the effect of genetic variations on APA patterns and shed light on future mechanistic understanding in APA regulation. Our code and data are available at https://github.com/lzx325/DeeReCT-APA-repo.

## Introduction

In eukaryotic cells, the termination of Pol II transcription involves 3’-end cleavage followed by addition of a poly(A) tail, a process termed as “polyadenylation”. Often, one gene could have multiple polyadenylation sites (PAS). The so-called alternative polyadenylation (APA) could generate from the same gene locus different transcript isoforms with different 3’-UTRs and sometimes even different protein coding sequences. The diverse 3’-UTRs generated by APA may contain different sets of *cis*-regulatory elements, thereby modulating the mRNA stability [1–3], translation [4], subcellular localization of mRNAs [5–7], or even the subcellular localization and function of the encoded proteins [8]. Importantly, it has been shown that dysregulation of APA could result in various human diseases [9–12].

APA is regulated by the interaction between *cis*-elements located in the vicinity of PAS and the associated *trans*-factors [13]. The most well-known *cis*-element that defines a PAS is the hexamer AAUAAA and its variants located 15-30nt upstream of the cleavage site, which is directly recognized by the cleavage and polyadenylation specificity factor (CPSF) components: CPSF30 and WDR33 [14]. Other auxiliary *cis-* elements located upstream or downstream of the cleavage site include upstream UGUA motifs bound by the cleavage factor Im (CFIm) and downstream U-rich or GU-rich elements targeted by the cleavage stimulation factor (CstF) [14]. The usage of individual PAS for a multi-PAS gene depends on how efficiently each alternative PAS is recognized by these 3’ end processing machineries, which is further regulated by additional RNA binding proteins (RBPs) that could enhance or repress the usage of distinct PAS signals through binding in their proximity. In addition, the usage of alternative PAS is mutually exclusive. In particular, once an upstream PAS is utilized, all the downstream ones would have no chance to be used no matter how strong their PAS signals are. Therefore, proximal PAS, which are transcribed first, have positional advantage over the distal competing PAS [15]. Indeed, it has been observed that the terminal PAS more often contain the canonical AAUAAA hexamer, which is considered to have higher affinity than the other variants, which possibly compensates for their positional disadvantage [16].

There has been a long-standing interest in predicting PAS based on genomic sequences using purely computational approaches. The so-called “PAS recognition problem” aims to discriminate between nucleotide sequences that contain a PAS and those do not. A variety of hand-crafted features have been proposed and statistical learning algorithms, *e*.*g*., random forest (RF), support vector machines (SVM) and hidden Markov models (HMM), are then applied on these features to solve the binary classification problem [17–19]. Very recently researchers started investigating the “PAS quantification problem”, which aims to predict a score that represents the strength of a PAS [20, 21]. This is much more difficult than the recognition one.

Recent developments in deep learning have made great improvements on many tasks [22]. With remarkable success, it has also been applied to bioinformatics tasks such as protein-DNA binding [23], RNA splicing pattern prediction [24], enzyme function prediction [25, 26], Nanopore sequencing [27, 28], and promoter prediction [29]. Deep learning is favored due to its automatic feature extraction ability and good scalability with large amount of data. As for polyadenylation prediction, deep learning models have been applied on the PAS recognition problem and they outperformed existing feature-based methods by a large margin [30]. Recently, deep learning models have also been applied on the PAS quantification problem, where Polyadenylation Code [20] was developed to predict the stronger one from a given pair of two competing PAS. Very recently, another model, DeepPASTA [21] has been proposed. DeepPASTA contains four different modules that deal with both the PAS recognition problem and PAS quantification problem. Similar as Polyadenylation Code, DeepPASTA also casts the PAS quantification problem into a pairwise comparison task.

In this paper, we propose a novel deep learning method, DeeReCT-APA (Deep Regulatory Code and Tools for Alternative Polyadenylation), for the PAS quantification problem. DeeReCT-APA can quantitatively predict the usage of all the competing PAS from a same gene simultaneously, regardless of the number of PAS. The model is trained and evaluated based on the dataset from a previous study [31], which consists of a genome-wide PAS measurement of two different mouse strains (C57BL/6J (BL) and SPRET/EiJ (SP)), and their F1 hybrid. After training our model on the dataset, we comprehensively evaluate our model based on a number of criteria. We demonstrate the necessity of modeling the competition among multiple PAS simultaneously. Finally, we show that our model can predict the effect of genetic variations on APA patterns, visualize APA regulatory motifs and potentially facilitate the mechanistic understanding of APA regulation.

## Methods

### Description of DeeReCT-APA architecture

The DeeReCT-APA method is based on a deep learning architecture that contains a set of neural network models composed of base networks (Base-Net, one for each competing PAS) and upper-level interaction layers. Each base network takes a 455nt long genomic DNA sequence centered around one competing PAS cleavage site as input and gives as output a vector which can be interpreted as the distilled features of that sequence. There are two types of base networks in our design, based on: (1) hand-engineered feature extractor and (2) convolutional neural networks (CNN). The output of the lower-level base network is then passed to the upper-level interaction layers, which computationally model the process of choosing competing PAS. The interaction layers of DeeReCT-APA are based on Long Short Term Memory Networks (LSTM) [32], which have achieved remarkable success in natural language processing and can naturally handle sentences with an arbitrary length, therefore suitable for handling any number of alternative PAS from a same gene locus. The interaction layers then output the percentage values of all the competing PAS of the gene. The architecture is illustrated in **Figure 1**. The design of each part of the network is further explained in the following subsections.

**Figure 1.**
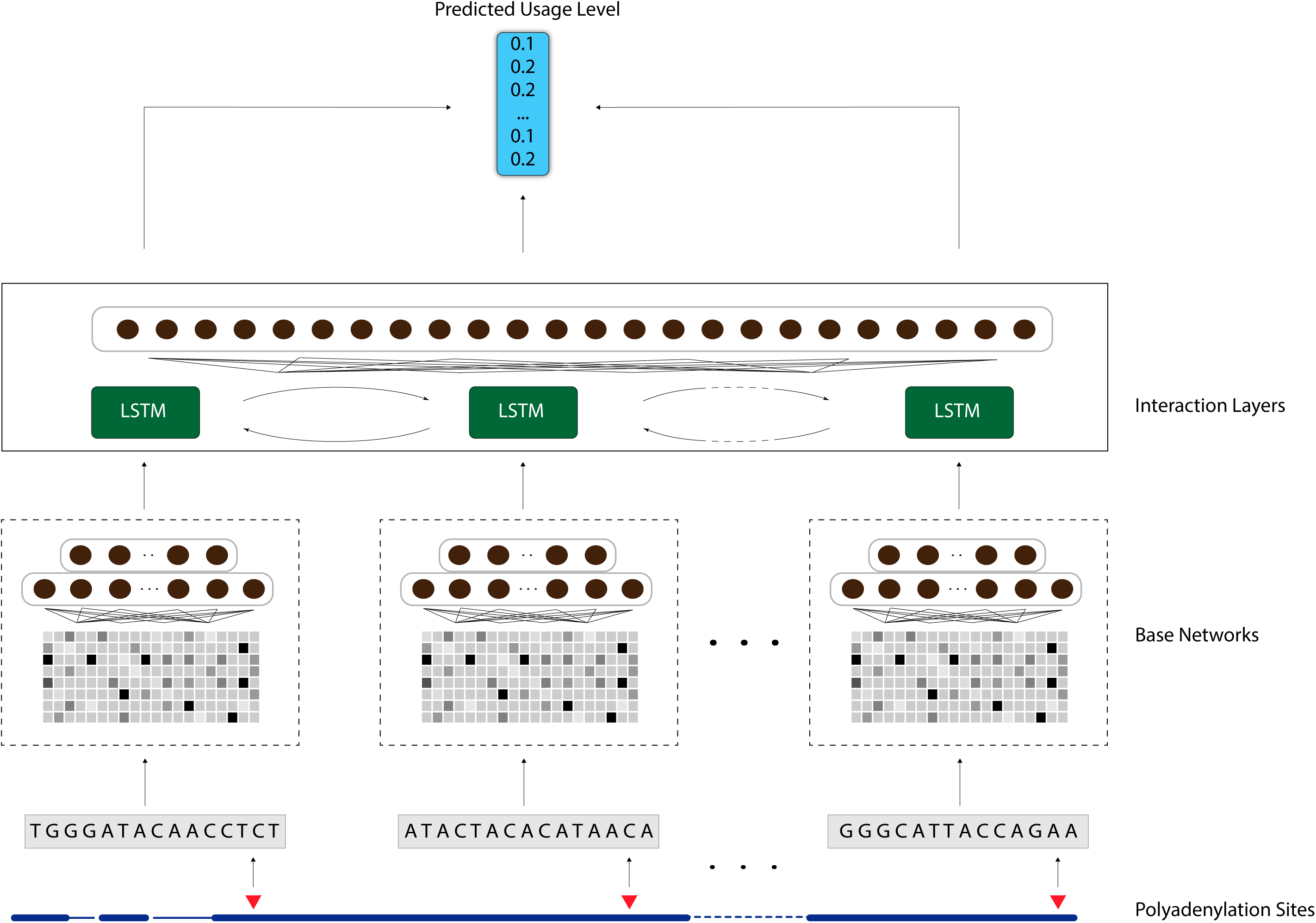
Illustration of the DeeReCT-APA architecture (Using BiLSTM as interaction layer)

We use three different base network designs: deep neural network architectures based on a single 1D convolution layer (Single-Conv-Net), multiple 1D convolution layers (Multi-Conv-Net) and a handcrafted feature extractor with fully-connected layers (Feature-Net). Single-Conv-Net and Multi-Conv-Net are two convolutional neural network (CNN) structures for Base-Net. The Single-Conv-Net consists of only one layer of the 1D convolutional layer and takes directly the one-hot encoded sequences as input. The convolutional layer has a number of convolution filters which become automatically-learned feature extractors after training. A rectified linear unit (ReLU) is used as the activation function. The max-pooling operation after that allows only values from highly-activated neurons to pass to the upper fully-connected layers. The three operations: convolution, ReLU and max-pooling form a convolution block. While the Single-Conv-Net uses one convolution block, the Multi-Conv-Net uses two convolution blocks before fully-connected layers. The increased depth of the network makes it possible for the network to learn more complex representations. The structures of Single-Conv-Net and Multi-Conv-Net are shown in **Figure 2A** and **Figure 2B**, respectively.

**Figure 2.**
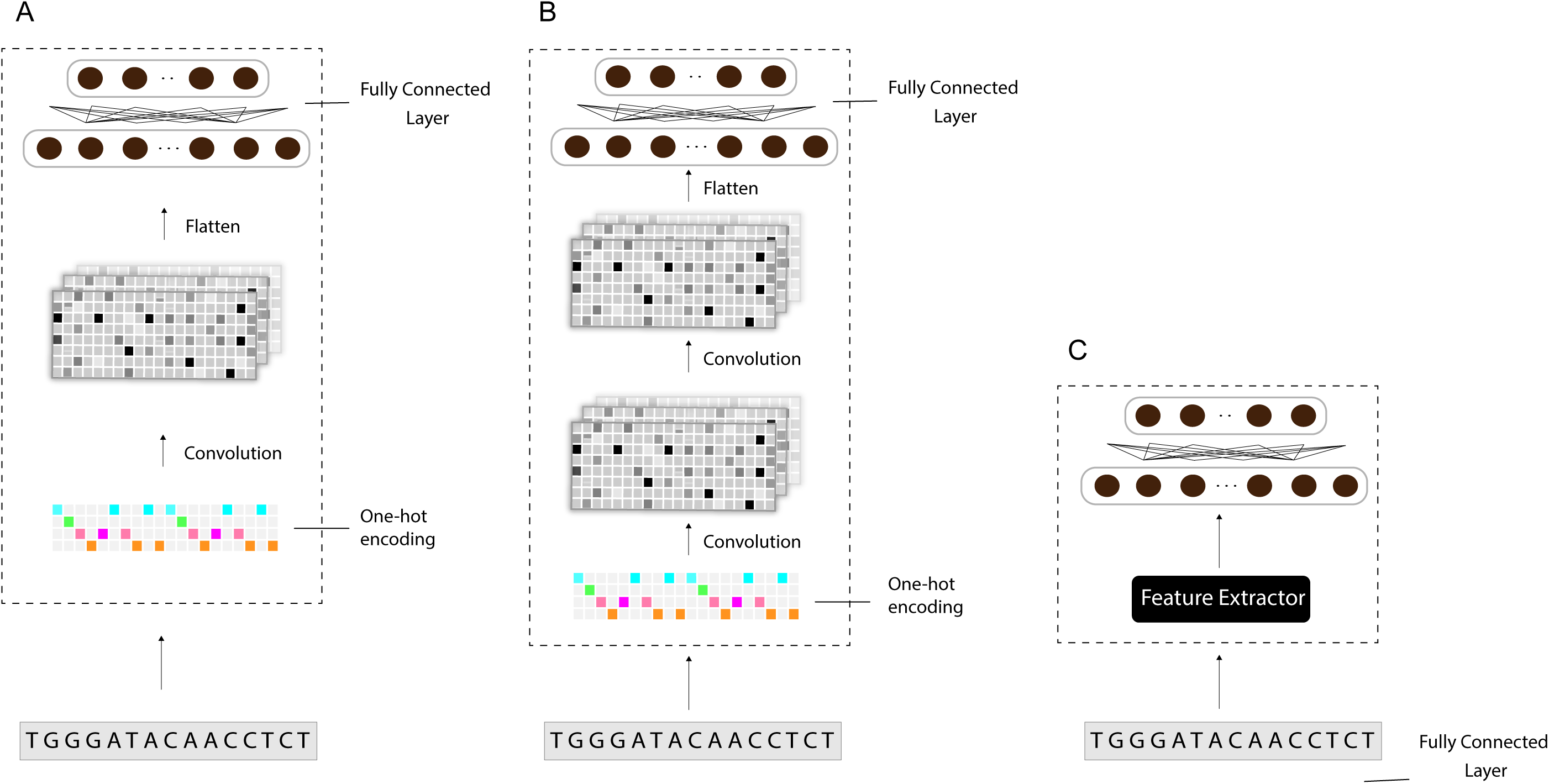
Three designs of Base-Net. All three of them output a feature vector that represents distilled features of the input sequence. **A**. Single-Conv-Net uses a single convolution layer for feature extraction. **B**. Multi-Conv-Net uses multiple convolution layers for feature extraction. **C**. Feature-Net contains a hand-crafted feature extractor before being processed by fully-connected layers.

As a comparison, we also design a base network that works with hand engineered features which we call Feature-Net. The Feature-Net only consists of multiple fully-connected layers and takes as input multiple types of features extracted from the sequences of interest. The features, described in [20], include polyadenylation signals, auxiliary upstream elements, core upstream elements, core downstream elements, auxiliary downstream elements [33], RNA-binding protein motifs, as well as 1-mer, 2-mer, 3-mer, and 4-mer features (detailed in Supplementary Materials Section S1 and Supplementary Table S1). Each feature value corresponds to the occurrence of each motif. The extracted features are then z-score normalized. The architecture is illustrated in **Figure 2C**.

### Design of the interaction layers

The utilization of alternative PAS is intrinsically competitive. On the one hand, as a multi-PAS gene is transcribed, any one of its PAS along the already transcribed region is possible to be used. But if one of them has already been used, it will make other PAS impossible to be chosen. On the other hand, given that the same polyadenylation machinery is used by all the alternative PAS, such competition of resources also contributes to the competitiveness of this process. However, previous work in polyadenylation usage prediction did not take this important point into account [20, 21]. Both existing models, Polyadenylation Code and DeepPASTA (tissue-specific relatively dominant poly(A) sites prediction model, Section 2.3 in [21]) can only take in two PAS at a time, ignoring the competition with others. Here, to overcome this limitation, we consider all the competing PAS at the same time and take as input all the PAS in a gene simultaneously into our model, then jointly predict the usage levels of all of them.

To fulfil this, we design the interaction layers above the base networks to model the interaction between different PAS. In neural networks, the most common way to model interactions among inputs is to introduce a recurrent neural network (RNN) layer, which can capture the interdependencies among inputs corresponding to each time step. We decide to choose the LSTM [32] as the foundation of interaction layers. LSTM is a type of RNN that has hidden memory cells which are able to remember a state for an arbitrary length of time steps, making it one of the most popular RNNs. To fit into the PAS usage level prediction task, each time step of LSTM corresponds to one PAS, at which the LSTM takes the extracted features of that PAS from the lower-level base network. As there is both influence from upstream PAS to downstream PAS and vice versa, we decide to use a bidirectional LSTM (BiLSTM), in which one LSTM’s time step goes from upstream PAS to downstream one and the other from downstream to upstream. The outputs of the two LSTMs at the same PAS are then concatenated and sent to the upper fully-connected layer. The fully-connected layer transforms the LSTM output to a scalar value representing the log-probability of that PAS to be used. After the log-probabilities of all competing PAS pass through a final SoftMax layer, they are transformed to properly normalized percentage scores, which sum up to one, representing their probability of being chosen. The detailed architecture is shown in Figure 1. We point out that, although DeepPASTA also contains a BiLSTM component, their BiLSTM layer is to process the sequence of one of the two competing PAS that are given as input. The time steps of the BiLSTM correspond to different positions in one particular sequence rather than to different PAS, and therefore the BiLSTM is not to model the interactions between different PAS, which is clearly different from the design in DeeReCT-APA.

As mentioned above, the aim of our model is to take all PAS of a gene as a whole and try to predict the usage level of each PAS as accurate as possible. Therefore, at one time, we must take all PAS in a gene as input. Considering that the number of PAS within a gene is not a constant, we design our model to take inputs of a variable length. Since most genes have a small number of PAS, we choose not to pad all the genes with dummy PAS to make them of the same length, otherwise it will be highly inefficient. Instead, we design the interaction layers in a way that it can take an arbitrary number of Base-Nets.

We further design two experiments for ablation study of DeeReCT-APA’s BiLSTM interaction layer. The first is to remove the BiLSTM layer and only keep the fully-connected layer and the SoftMax layer. In this scenario, the network still considers all PAS of a gene simultaneously, but with a non-RNN interaction layer. The second is to remove the interaction layer altogether and use comparison-based training (like in Polyadenylation Code) to train a Base-Net. We show their performance separately in the “Overall Performance” section.

### A genome-wide PAS quantification dataset derived from fibroblast cells of C57BL/6J (BL) and SPRET/EiJ (SP) mouse and their F1 hybrid

A genome-wide PAS quantification dataset derived from fibroblast cells of C57BL/6J (BL) and SPRET/EiJ (SP), as well as their F1 hybrid is obtained from the previous study [31]. In the F1 cells, the two alleles have the same *trans* environment and the PAS usage difference between two alleles can only be due to the sequence variants between their genome sequences, making it a valuable system for APA *cis*-regulation study. Apart from APA, this kind of systems have also been used in the study of alternative splicing and translational regulation [34, 35].

The detailed description of the sequencing protocol and data analysis procedure can be found in [31]. As a brief summary, the study uses fibroblast cell lines from BL, SP and their F1 hybrids. The total RNA is extracted from fibroblast cells of BL and SP undergoes 3’-Region Extraction and Deep Sequencing (3’READS) [16] to build a good PAS reference of the two strains. The 3’-mRNA sequencing is then performed in all three cell lines to quantify those PAS in the reference. In the F1 hybrid cell, reads are assigned to BL and SP alleles according to their strain specific SNPs. The PAS usage values are then computed by counting the sequencing reads assigned to each PAS. The sequence centering around each PAS cleavage site (448nt in total) is extracted and undergoes feature extraction or one-hot encoding before training the model. The extracted features are then inputted to Feature-Net, while the one-hot encoded sequences are inputted to Single-Conv-Net and Multi-Conv-Net.

### Training and evaluation of the model

We train the DeeReCT-APA models based on the parental BL/SP PAS usage level dataset. For F1 hybrid data, however, we choose to start from the pre-trained parental model (which we use either the BL parental model or the SP parental model and the results are shown separately) and fine-tune the model on the F1 dataset. This is because, due to the read assignment problem, the usage of many PAS in F1 cannot be unambiguously characterized by 3’-mRNA sequencing [31]. As a result, the F1 dataset does not contain enough number of PAS to train our model from scratch. At the training stage, genes are randomly selected from the training set and the sequences of their PAS flanking regions are fed into the network. Each sequence of PAS in a gene passes through one Base-Net. The parameters of the Base-Net that are responsible for each PAS are all shared. The Base-Net then each outputs a vector representing distilled features for each PAS, which is then sent to the interaction layers. The interaction layers generate a percentage score of each PAS of this gene. Cross-entropy loss between the predicted usage and the actual usage is used as the training target. During back-propagation, the gradients are back-propagated through the passage originated from each PAS. As the model parameters are shared between base networks, the gradients are then summed up to update the model parameters. We use several techniques to reduce overfitting: (1) Weight decay is applied on weight parameters of CNN and all fully-connected layers. (2) Dropout is applied on BiLSTM. (3) We stop training as soon as the mean absolute error of the predicted usage value does not improve on the validation set. (4) While fine-tuning the model on F1 dataset, we use a learning rate that is ∼100 times smaller than the one used when training from scratch.

The network is trained with the adaptive moment estimation (Adam) optimizer [36]. A detailed list of hyperparameters we used is specified in Supplementary Materials Section S2 and Supplementary Table S2. We construct the network using the PyTorch deep learning framework [37] and utilize one NVIDIA GeForce GTX 980 Ti as the GPU hardware platform.

To evaluate the performance of the model, we conduct a 5-fold cross validation at the gene level using all the genes in our dataset for each strain. That is, if a gene is selected as a training (testing) sample, all of its PAS are in the train (test) set. At each time, four folds are used for training and the remaining one is used for testing. To make a fair comparison with Polyadenylation Code and DeepPASTA in Section 3.1, we also train (fine-tune) the two models and optimize their model parameters on the parental and F1 datasets.

### Performance measures

To comprehensively evaluate DeeReCT-APA and compare it against baseline and state-of-the-art methods, we use the following performance measures.

#### Mean Absolute Error (MAE)

This metric is defined as the mean absolute error (MAE) of the usage prediction of each PAS, which is

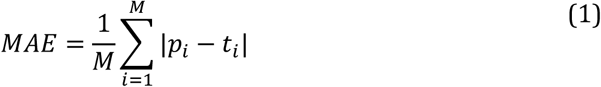

where *p*_*i*_ stands for the predicted usage, *t*_*i*_ stands for the experimentally determined ground truth usage for PAS i and M is the total number of PAS across all genes in the test set. This is the most intuitive way of measuring the performance of DeeReCT-APA. However, it is not applicable to Polyadenylation Code [20] or DeepPASTA [21] as they do not have quantitative outputs that can be interpreted as the PAS usage values. For the same reason, it is not applicable to DeeReCT-APA either, when its interaction layers are removed and use comparison-based training (Section “Design of the interaction layers”).

#### Comparison Accuracy

We here define the Pairwise Comparison Task. We enumerate all the pairs of PAS in a given gene and keep those pairs with PAS usage level difference greater than 5%. We then ask the model to predict which PAS in the pair is of the higher usage level. The accuracy is defined as,

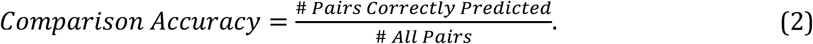

Note that the primary reason that we use this metric is to compare with Polyadenylation Code and DeepPASTA, as they were designed for predicting which one is stronger between the two competing PAS.

#### Highest Usage Prediction Accuracy

We here define the Highest Usage Prediction Task. This task aims to test the model’s ability of predicting which PAS is of the highest usage level in a single gene. We select all the genes which has its highest PAS usage level greater than its second highest one by at least 15% in the test set for evaluation. For DeeReCT-APA, the predicted usage in percentage is used for ranking the PAS. For Polyadenylation Code and DeepPASTA, as they do not provide a predicted value in percentage, the logit value before the SoftMax layer is used instead. The logit values, though not in the scale of real usage percentage values, can at least give a ranking of different PAS sites. The highest usage prediction accuracy is the percentage of genes whose highest-usage PAS are correctly predicted.

#### Averaged Spearman’s Correlation

We here define the Ranking Task. We convert the predicted usage levels by each model into a ranking of PAS sites in that gene. We then compute the Spearman’s correlation between the predicted ranking and ground truth ranking. The correlation values for all genes are then averaged together to give an aggregated score. In other words,

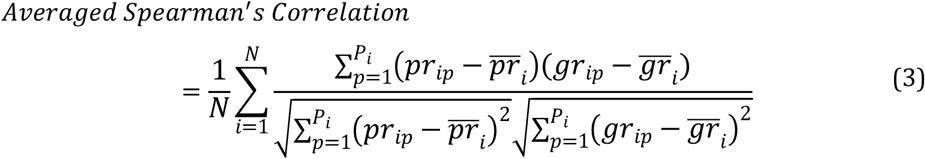

where *N* is the total number of genes, *P*_*i*_ is the number of PAS in gene *i, pr*_*ip*_is the predicted rank of PAS *p* in gene *i, gr*_*ip*_is the ground truth rank of PAS p in gene i, *pr*_*i*_ and *gr*_*i*_ are averaged predicted and ground truth ranks in gene i, respectively.

## Results

### Overall performance

First, to compare the performance of different Base-Net designs, we evaluated DeeReCT-APA with different Base-Nets: Feature-Net, Single-Conv-Net, and Multi-Conv-Net. As shown in Supplementary Table S3, both on the parental BL dataset and on the F1 dataset, DeeReCT-APA with Multi-Conv-Net performs the best, followed by that with Single-Conv-Net. This is expected, as deeper neural networks have higher representation learning capacity.

We then compared the performance of DeeReCT-APA with Multi-Conv-Net to Polyadenylation Code and DeepPASTA. As shown in **Table 1**, both on the parental BL dataset and on the F1 dataset, DeeReCT-APA with Multi-Conv-Net consistently performs the best across all four metrics. The standard deviation across 5-fold cross validation is higher in the F1 dataset than in the parental dataset, indicating the increased instability in F1 prediction which is probably due to the limited amount of F1 data. As we have a rather small dataset, a very complex model like DeepPASTA is prone to overfitting, which is probably the reason why it performs the worst here. Indeed, for the smaller F1 dataset, DeepPASTA lags even more behind other methods.

**Table 1.** Performance summary for the BL parental model and the F1 model.

| Model | Performance on Parental Dataset |  |  |  |
| --- | --- | --- | --- | --- |
|  | MAE* | Comparison Accuracy* | Highest Usage Prediction Accuracy* | Averaged Spearman's Correlation |
| DeeReCT-APA (Multi-Conv-Net) | **17.22% ± 0.3%** | **77.64% ± 0.4%** | **63.48% ± 0.9%** | **0.5140 ± 0.021** |
| Polyadenylation Code | N/A | 75.88% ± 0.8% | 59.82% ± 1.5% | 0.4673 ± 0.022 |
| DeepPASTA | N/A | 74.08% ± 1.1% | 58.78% ± 1.4% | 0.4394 ± 0.017 |

\*The values for a random predictor are 43.12%, 50.00% and 25.49% respectively. MAE, mean absolute error. Note that for MAE, it is the *lower* the better.

B
| Model | Performance on F1 Dataset |  |  |  |
| --- | --- | --- | --- | --- |
|  | MAE* | Comparison Accuracy* | Highest Usage Prediction Accuracy* | Averaged Spearman's Correlation |
| DeeReCT-APA (Multi-Conv-Net) | **17.80% ± 0.3%** | **77.14% ± 1.2%** | **64.52% ± 0.7%** | **0.4567 ± 0.009** |
| Polyadenylation Code | N/A | 74.20% ± 0.1% | 59.04% ± 0.9% | 0.4224 ± 0.014 |
| DeepPASTA | N/A | 70.14% ± 1.5% | 53.82% ± 1.7% | 0.3693 ± 0.018 |

\*The values for a random predictor are 40.96%, 50.00% and 28.56% respectively. MAE, mean absolute error. Note that for MAE, it is the *lower* the better.
*Note:* The parental model is trained from scratch and the F1 model is fine-tuned from the BL parental model. The table shows the performance of three models across four evaluation metrics. Results are shown in the mean±std format. The best performance is in bold. See Section “Overall performance” for details. A. Performance on the Parental Dataset (BL). B. Performance on the F1 Dataset (fine-tuned from parental BL model).

Similar results on the parental SP dataset and the performance of F1 model that is fine-tuned from the SP parental model are shown in Supplementary Materials Section S3 and Supplementary Table S4. Unless otherwise stated, the F1 model that we use in the remaining part of the paper is the one fine-tuned from the parental BL model and using the training set folds that do not include the gene or PAS to be tested.

Next, we show that, in terms of comparison accuracy, the improvement made by DeeReCT-APA is statistically significant, even though the performance improvement is not numerically substantial. For this purpose, we repeat the experiment for five times, with each of them having the dataset randomly split in a different way, and report the accuracy of DeeReCT-APA (Multi-Conv-Net), Polyadenylation Code, and DeepPASTA after 5-fold cross validation (Supplementary Materials Section S4 and Supplementary Table S5). The performance of three tools is then compared with p-value computed by t-test. As shown in Supplementary Table S5, indeed the improvement of DeeReCT-APA over the other two methods is statistically significant.

To demonstrate that the results of our comparison is independent of the datasets, we train and test DeeReCT-APA also on another dataset used in [20]. Since it consists of polyadenylation quantification data from multiple human tissues, we report the performance (comparison accuracy) of DeeReCT-APA for each tissue separately (Supplementary Materials Section S4 and Supplementary Table S6). The performance metrics of Polyadenylation Code and DeepPASTA is adapted from [20] and [21] accordingly. For 6 out of 8 tissues, DeeReCT-APA achieves higher accuracy than the other two methods.

We finally show through ablation study that the usage of BiLSTM interaction layer contributes to the performance of DeeReCT-APA. As shown in **Table 2**, we compare the performance of DeeReCT-APA with Multi-Conv-Net (1) without interaction layer, to (2) with interaction layer but without BiLSTM, and (3) with interaction layer and with BiLSTM (The detailed architectures are shown in Supplementary Figure S1). In terms of all metrics, both the usage of interaction layer and BiLSTM improve the performance. Although not numerically substantial, the improvements are in general statistically significant after performing a similar experiment as we have done earlier (Supplementary Table S7). The improvement of (2) over (1) (p=2.5e-6 for parental and p=1.1e-3 for F1) is more statistically significant than (3) over (2) (p=3.7e-3 for parental and p=9.9e-2 for F1) indicating that the majority of the performance gain of DeeReCT-APA comes from using the interaction layers and the simultaneous consideration of all PAS. This concludes that DeeReCT-APA, with an RNN interaction layer that considers all PAS of a gene at the same time, can achieve better performance on the PAS quantification task.

**Table 2.** The performance of DeeReCT-APA using different interaction layers.

| Model | Performance on Parental Dataset |  |  |  |
| --- | --- | --- | --- | --- |
|  | MAE* | Comparison Accuracy* | Highest Usage Prediction Accuracy* | Averaged Spearman's Correlation |
| DeeReCT-APA<br>(Multi-Conv-Net)<br>(No Interaction Layer) | - | 76.12% ± 0.5% | 60.02% ± 0.7% | 0.4988 ± 0.027 |
| DeeReCT-APA<br>(Multi-Conv-Net)<br>(w/o BiLSTM) | 17.54% ± 0.3% | 77.12% ± 0.5% | 61.73% ± 0.6% | 0.5007 ± 0.034 |
| DeeReCT-APA<br>(Multi-Conv-Net)<br>(BiLSTM) | **17.22% ± 0.3%** | **77.64% ± 0.4%** | **63.48% ± 0.9%** | **0.5140 ± 0.021** |

\* The values for a random predict or are 43.12%, 50.00% and 25.49% respectively. MAE, mean absolute error. Note that for MAE, it is the lower the better.

B
| Model | Performance on F1 Dataset |  |  |  |
| --- | --- | --- | --- | --- |
|  | MAE* | Comparison Accuracy* | Highest Usage Prediction Accuracy* | Averaged Spearman's Correlation |
| DeeReCT-APA<br>(Multi-Conv-Net)<br>(No Interaction Layer) | - | 76.28% ± 1.1% | 61.72% ± 0.8% | 0.4337 ± 0.019 |
| DeeReCT-APA<br>(Multi-Conv-Net)<br>(w/o BiLSTM) | 18.03% ± 0.2% | 76.77% ± 1.0% | 63.44% ± 0.3% | 0.4751 ± 0.011 |
| DeeReCT-APA<br>(Multi-Conv-Net)<br>(BiLSTM) | **17.80% ± 0.4%** | **77.14% ± 1.2%** | **64.52% ± 0.7%** | **0.4957 ± 0.009** |

\*The values for a random predictor are 40.96%, 50.00% and 28.56% respectively. MAE, mean absolute error. Note that for MAE, it is the lower the better.
*Note:* The table shows the performance of DeeReCT-APA with different interaction layers. Note that for DeeReCT-APA without interaction layer, the model is trained based on comparison and its output cannot
be interpreted as a percentage score. Therefore, like for Polyadenylation Code and DeepPASTA earlier, we do not report its MAE value. A. Performance on the Parental Dataset (BL). B. Performance on the F1 Dataset (fine-tuned from parental BL model).

### Benefits of modelling all PAS jointly—one example

To illustrate DeeReCT-APA’s ability of modeling all PAS of a gene jointly, we use the gene *Srr* (Ensembl Gene ID: ENSMUSG00000001323) as an example. As shown in **Figure 3A**, the gene *Srr* use four different PAS, whereas **Figure 3B, 3C, 3D** shows the ground truth usage level, the prediction of DeeReCT-APA with Multi-Conv-Net and Polyadenylation Code, in the F1 hybrid cell for those four PAS, for both its BL allele (blue bars) and SP allele (green bars), respectively. As before, the logits values before the SoftMax layer of Polyadenylation Code are used as surrogates for predicted usage values (and therefore not in the range from 0 to 1). As shown in **Figure 3**, the prediction of DeeReCT-APA is much more consistent with the ground truth than that of Polyadenylation Code and the relative magnitude between the BL allele and SP allele for the prediction of DeeReCT-APA is correct for all four PAS. In comparison, Polyadenylation Code model predicted PAS 4 in the BL allele to be of slightly *higher* usage than the one in the SP allele whereas both in ground truth and the prediction made by DeeReCT-APA, the reverse is true. We hypothesize in this case that the genetic variants between the BL allele and SP allele in the sequences flanking PAS 4 alone might make the BL allele a *stronger* PAS than the SP allele because Polyadenylation Code only considers which one between the two is stronger and predicts the strength of a PAS solely by its own sequence, without considering those of the others. However, when simultaneously considering genetic variations in PAS 1, PAS 2, and PAS 3, which probably have *stronger* effects, the usage of PAS 4 becomes *lower* in BL than in SP.

**Figure 3.**
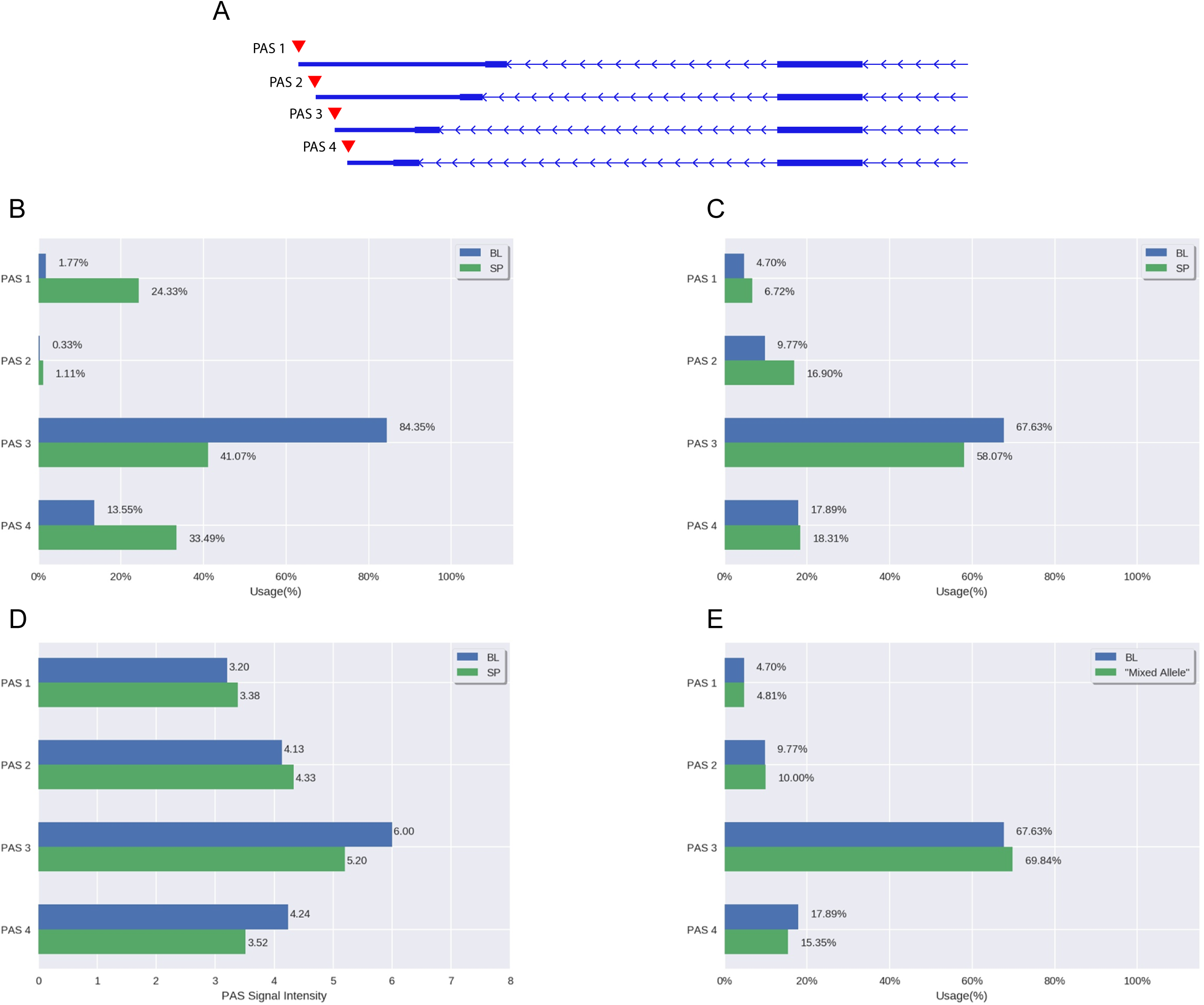
Prediction of gene *Srr*. This shows one example of the benefit of modelling all PAS jointly. Each panel shows the predicted or ground truth usage of each of its four PAS: **A**. PAS of gene *Srr*. **B**. Ground Truth. **C**. DeeReCT-APA’s (Multi-Conv-Net) prediction. **D**. Polyadenylation Code’s prediction. **E**. DeeReCT-APA’s (Multi-Conv-Net) prediction of “mixed allele”. The prediction of DeeReCT-APA is much more consistent with ground truth compared to Polyadenylation Code. Especially for PAS 4, DeeReCT-APA predicts the one of BL allele to be of lower usage than the one of SP allele which is consistent with ground truth. Polyadenylation Code, on the contrary, predicts the opposite. In Panel **E**, by making prediction of the “Mixed Allele”, we demonstrated that the increased usage of PAS in SP allele is probably due to the concerted effects of the other three PAS.

To test our hypothesis, we design an *in-silico* experiment by constructing a hypothetical allele of gene *Srr* (hereafter referred to as “mixed allele”) that has the BL sequence of PAS 1, PAS 2, and PAS 3, and SP sequence of PAS 4. We then ask the DeeReCT-APA model to predict the usage level of each PAS in the “mixed allele”, where the usage differences between the BL allele and the “mixed allele” should then be purely due to the sequence variants in PAS 4 because the two alleles are exactly the same on the other PAS. As shown in **Figure 3E**, consistent with our hypothesis, the usage level of PAS 4 in the BL allele is indeed *higher* than that in the “mixed allele”. This example nicely demonstrates the benefit of jointly modeling all the PAS in a gene simultaneously.

### Allelic difference in PAS usage between BL and SP

One primary goal of developing DeeReCT-APA is to determine the effect of sequence variants on APA patterns. The F1 hybrid system we choose here is ideal to test how well such a goal is achieved, since in the F1 cells, the allelic difference in PAS usage can only be due to the sequence variants between their genome sequences.

Figure 4 shows two examples: gene *Zfp709* (Ensembl Gene ID:ENSMUSG00000056019) and *Lpar2* (Ensembl Gene ID: ENSMUSG00000031861), where previous analysis demonstrated that in the distal PAS of gene *Zfp709*, a substitution (from A to T) in the SP allele relative to the BL allele disrupted the PAS signal (ATTAAA to TTTAAA) (**Figure 4A**); in the distal PAS of gene *Lpar2*, a substitution (from A to G) in the SP allele relative to the BL allele disrupted another PAS signal (AATAAA to AATAAG) (**Figure 4B**), causing both of them to be of lower usage in the SP allele than in the BL allele.

To check whether our model could be used to identify the effects of these variants, we plot a “mutation map” for the two genes. In brief, for each gene, given the sequence around the most distal PAS (suppose it is of length L), we generate 3L “mutated sequences”. Each one of the 3L sequences has exactly one nucleotide mutated from the original sequence. These 3L sequences are then fed into the model along with other PAS sequences from that gene and the model then predicts usage for all sites and for each of the 3L sequences, separately. The predicted usage values of the original sequence are then subtracted from each of the 3L predictions and plotted in a heatmap, the “mutation map”.

**Figure 4.**
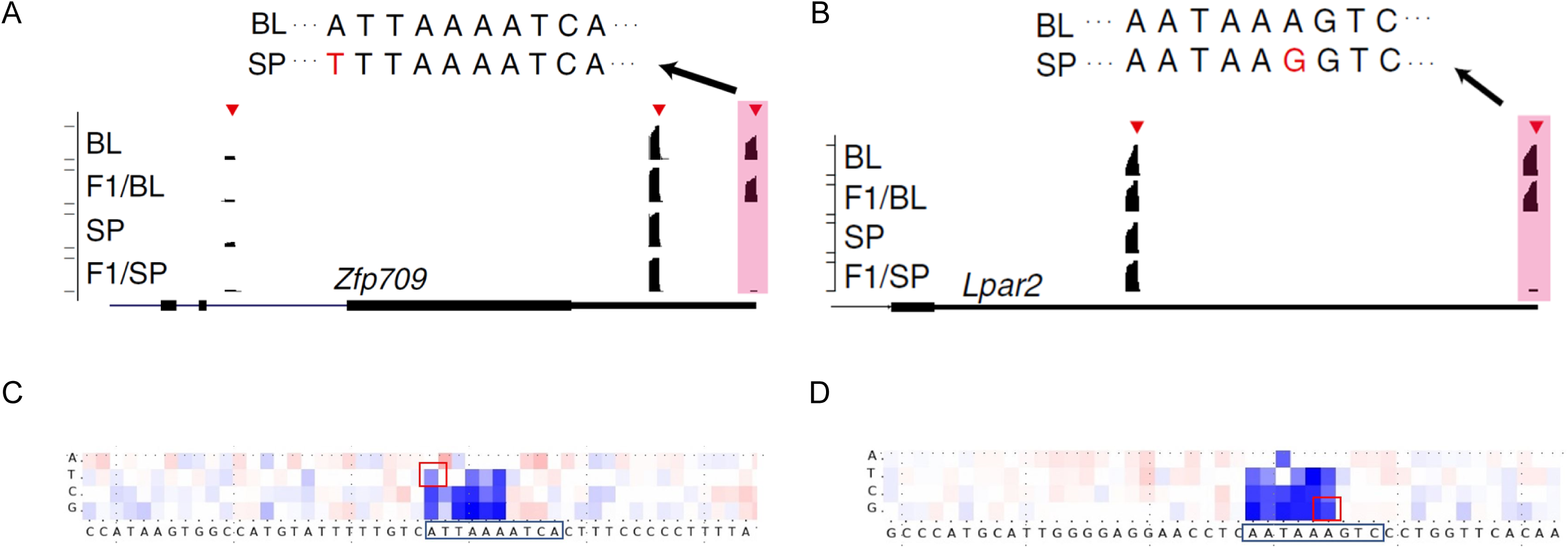
Previous experimental findings and mutation map of gene *Zfp709* and *Lpar2*. Mutation map is consistent with previous experimental findings on two genes, *Zfp709* (**A** & **C**) and *Lpar2* (**B** & **D**). Sequencing read coverage graphs (**A** & **B**) are adapted from Figure 4H of [31]. The identified PAS are marked by red triangles on top of the sequencing read coverage (black coverage graph). The sequence variants of the PAS shaded in pink between BL and SP strains are shown on the top. The BL mutation map (**C** & **D**) of the BL distal PAS sequence shows the effect of BL distal sequence mutation on the usage of distal sites. The SP gene *Zfp709* and *Lpar2* can be viewed as undergoing a substitution relative to BL. The four heatmap entries above each letter of the sequence (**C** & **D**, bottom) show the relative change of usage level when the nucleotide at that position is substituted with the nucleotide of the corresponding row. Darker red indicates greater increase in usage and darker blue indicates more decrease in usage. The entries that correspond to the genetic variants between BL and SP in **A** & **B** are marked by red squares.

As shown in **Figure 4C** and **Figure 4D**, the heatmap entries that correspond to the sequence variants between BL and SP is consistent with experimental findings from [31] (**Figure 4A and Figure 4B**). In addition, the mutation maps can also show the predicted effect of sequence variants other than those between BL and SP, giving an overview of the effects from all potential mutations.

Obviously, the two examples described above involved sequence variants disrupting PAS signals, which makes the prediction relatively trivial. To check whether our model could be used for the variants with more subtle effect, we choose a third example, gene *Alg10b*. Previous experiments showed that the usage of the most distal PAS of its BL allele is higher than its SP allele (**Figure 5A**). Using reporter assays (**Figure 5B**), it has been demonstrated that [31] an insertion of UUUU in the SP allele relative to the BL allele contributes to this reduction in usage (**Figure 5C**). To check whether DeeReCT-APA could reveal such effects, we also construct the same four *in silico* sequences as in [31] : BL, SP, BL2SP, and SP2BL. Together with other PAS of gene *Alg10b*, the four sequences are feed to the DeeReCT-APA model, separately. As shown in **Figure 5D**, comparing BL with BL2SP and SP with SP2BL, our model is able to reveal the negative effect of poly(U) tract.

**Figure 5.**
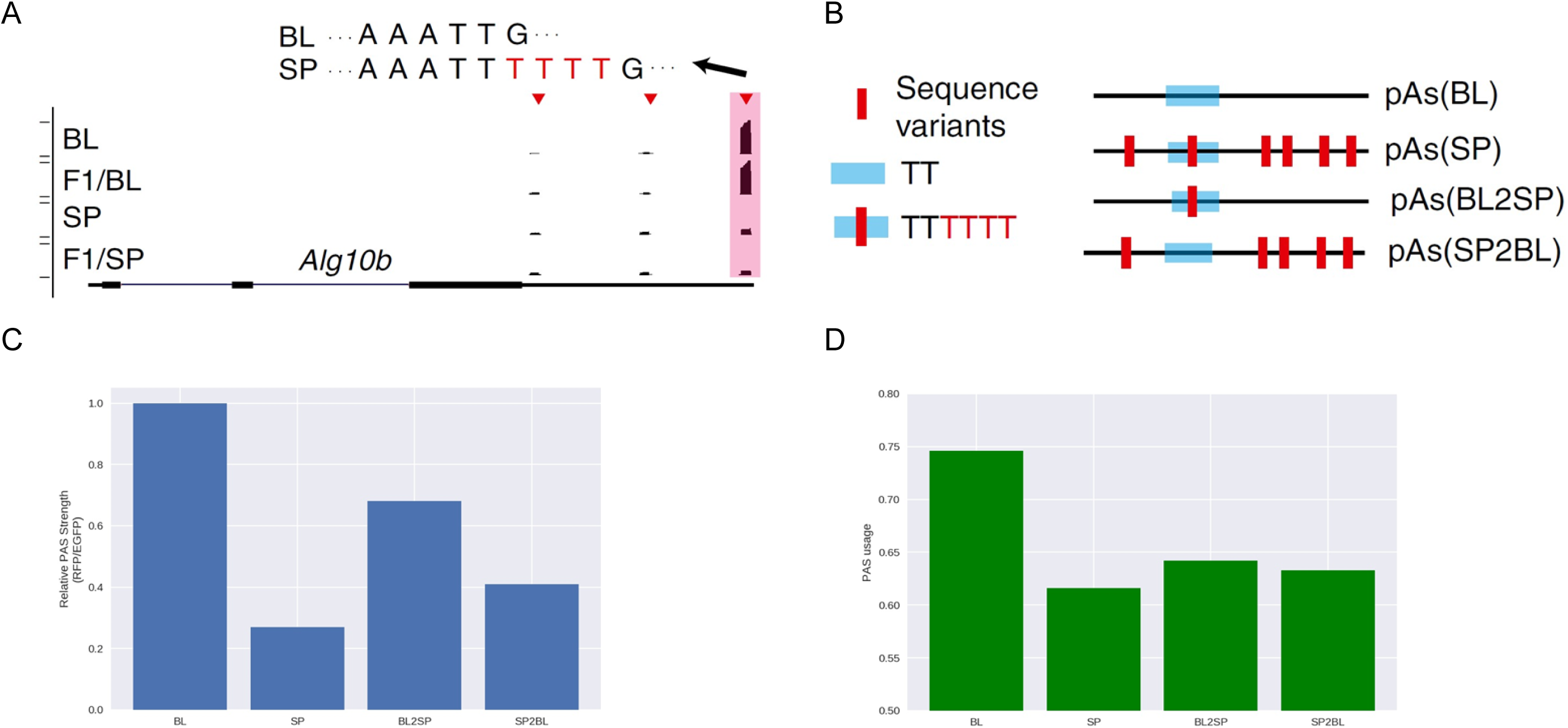
Previous experimental findings and DeeReCT-APA’s prediction of gene *Alg10b*. *In silico* prediction for the *Alg10b* PAS reporter is consistent with previous experimental findings. Similar to Figure 4A, the sequencing read coverage graph and the sequence variants are shown in **A**. The red triangles mark the identified PAS sites. The structures of PAS reporter constructs are shown in **B**, where “BL” is the original BL version of the most distal PAS, “SP” is the original SP version, “BL2SP” is the BL sequence only inserted with TTTT at the corresponding location and “SP2BL” is the SP sequence only deleted with TTTT at the corresponding location. The experimental results from PAS reporter assay for the four reporters are shown in **C**. and their *in silico* predictions are shown in **D**. Considering the *in silico* prediction pairs, BL & BL2SP and SP & SP2BL, it is clear that DeeReCT-APA is able to identify the negative modulation of PAS usage by the poly(U) tract. Figure (**A, B** & **C**) are adapted from Figure 4H of [31]. See text for more details.

To globally evaluate the performance of DeeReCT-APA on predicting the allelic difference in PAS usage, we compare the predicted allelic difference versus experimentally measured allelic difference in a genome-wide manner (**Figure 6A**). As a baseline control, we do the same for the prediction made by the Polyadenylation Code where logit values before SoftMax are again used as surrogates for the predicted allelic difference in PAS usage (**Figure 6B**). Here, the F1 model fine-tuned from the BL parental model is used. Similar results of the F1 model fine-tuned from the SP parental model are shown in Supplementary Materials Section S3 and Supplementary Figure S2. It is worth noting that this is a very challenging task because the training data do not well represent the complete landscape of genetic mutations. That is, the BL dataset only contains invariant sequences from different PAS, and the F1 dataset contains a limited number of genetic variants.

**Figure 6.**
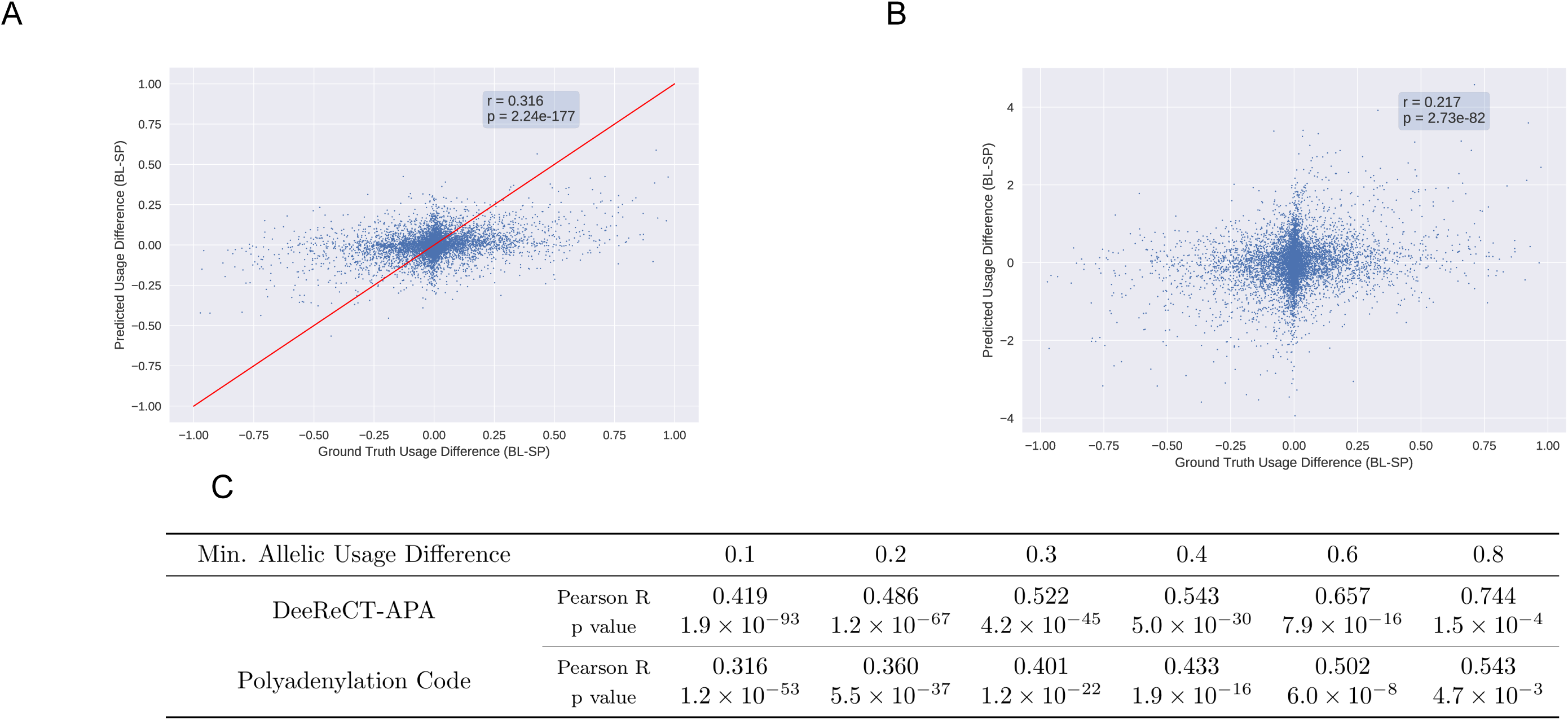
Comparison of the allelic usage difference predicted by DeeReCT-APA and Polyadenylation Code. F1 model fine-tuned from BL parental model is used. **A. B**. The horizontal axis is the ground truth allelic usage difference (BL usage minus the SP usage). The vertical axis shows the predicted allelic usage difference. The red line shows the perfect prediction. In terms of Person correlation, DeeReCT-APA shows better correlation than Polyadenylation Code. **C**. Pearson correlations (and their p-values) between two quantities at different minimum allelic usage difference are shown in the table below. The prediction of DeeReCT-APA still has better correlation than Polyadenylation Code when the dataset is filtered at different thresholds.

We then compute the Pearson correlation between the experimentally measured allelic usage difference and the ones predicted by the two models. Clearly, DeeReCT-APA outperforms Polyadenylation Code. We further evaluate the Pearson correlation values using six subsets of the test set, each filtering out PAS with allelic usage difference less than 0.1, 0.2, 0.3, 0.4, 0.6, 0.8, respectively (**Figure 6, Panel C**). When the allelic usage difference is small, their relative magnitudes are more ambiguous and the experimental measurement of their allelic usage difference (used here as ground truth) are less confident. Indeed, with the increasing allelic difference, the prediction accuracy increased for both DeeReCT-APA and Polyadenylation Code. Importantly, in all these groups, DeeReCT-APA shows consistently better performance.

### Visualization of convolutional filters

To show the knowledge learned by the convolutional filters of DeeReCT-APA, we follow a similar procedure as in [36] to visualize the convolutional filters of the model. The aim of visualization is to reveal the important subsequences around polyadenylation sites that activate a specific convolutional filter. In contrast to [38], in which the researchers only used sequences in the test set for visualization, we use all sequences in the train and test dataset of F1 for visualization due to the smaller size of our dataset. In visualization, neither the model parameters nor the hyperparameters are tuned on the test set, our usage of test set for visualization is therefore legitimate. For all the learned filters in layer 1, we convolve them with all the sequences in the above dataset, and for each sequence, its subsequence (having the same size as the filters) with the highest activation on that filter is extracted and accumulated in a position frequency matrix (PFM). The PFM is then ready for visualization as the knowledge learned by that specific filter. For layer 2 convolutional filters, as they do not convolve with raw sequences during training and testing, directly convolving it with the sequences in the dataset as we did for layer 1 would be undesirable. Instead, the layer 2 activations are calculated by a partial forward pass in the network and the subsequences of the input sequences in the receptive field of the maximally-activated neuron is extracted and accumulated in a PFM.

As shown in **Figure 7A** and **7B**, DeeReCT-APA is able to identify the two strongest PAS hexmer, AUUAAA and AAUAAA [31]. In addition, one of the layer 2 convolutional filters is able to recognize the pattern of a mouse specific PAS hexamer UUUAAA [30] (**Figure 7C**). Furthermore, a Poly-U island motif previously reported in [38] could also be revealed by DeeReCT-APA (**Figure 7D**). A complete visualization of all the 40 filters in layer 1 and 40 filters in layer 2 is shown in Supplementary Figure S3 and Supplementary Figure S4.

**Figure 7.**
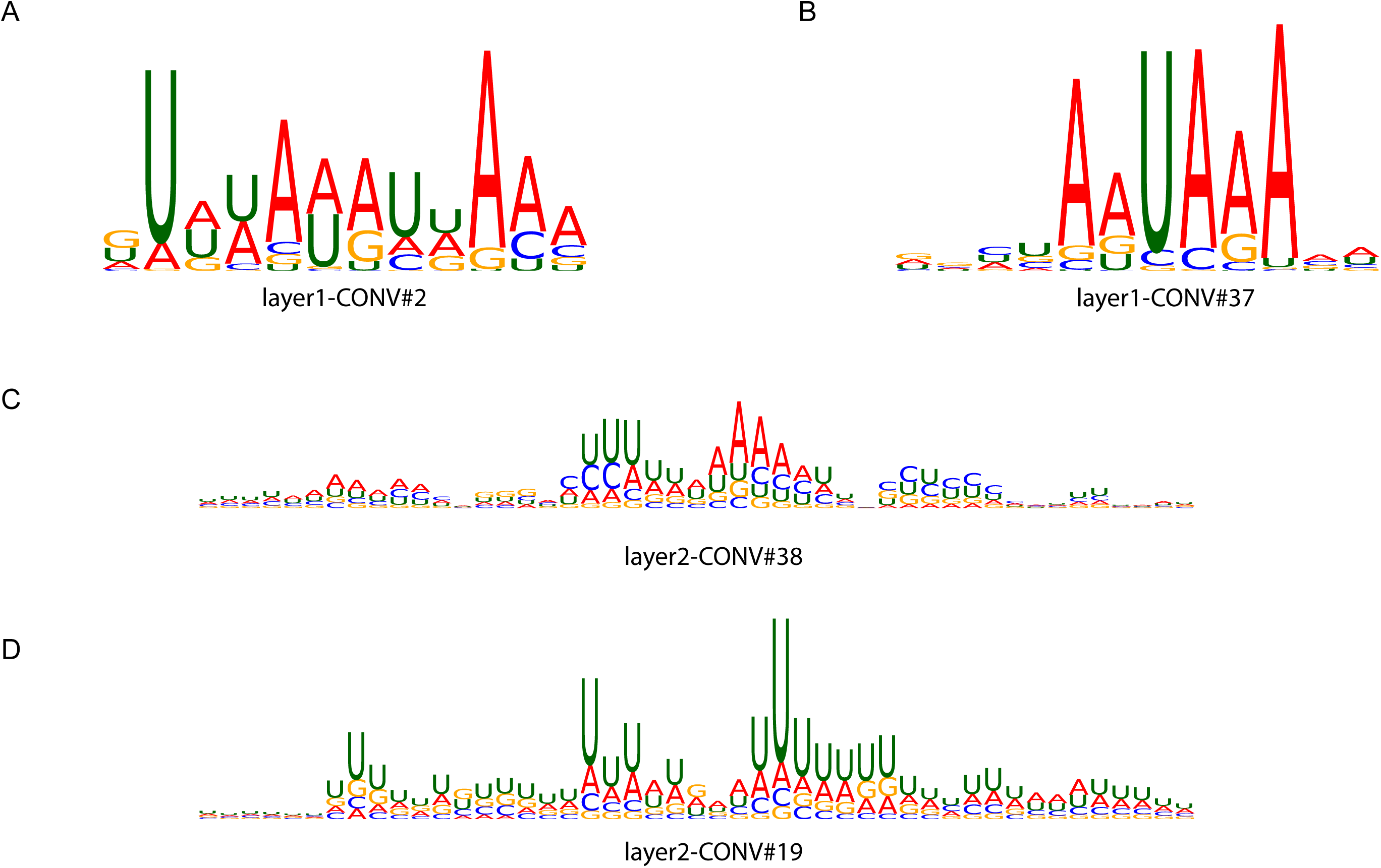
Visualization of learned convolutional filters in DeeReCT-APA. Some visualization examples of the learned convolutional filters of DeeReCT-APA. **A. B**. The most common polyadenylation motifs AUUAAA and AAUAAA are learned in layer 1 convolutional filter #2 and #37, respectively. **C**. Visualization of a layer 2 filter, #38 shows a mouse specific polyadenylation motif UUUAAA. **D**. Layer 2 filter #19 shows the Poly-U islands on polyadenylation. Note that the layer 2 filter visualization PFMs are wider than the layer 2 filter (12nt) because the receptive field of neurons in a deeper layer is in general greater than their corresponding filter width.

## Discussion and conclusion

In this study, we made the first attempt to simultaneously predict the usage of all competing PAS within a gene. Our method incorporates both sequence-specific information through automatic feature extraction by CNN and multiple PAS competition through interaction modeling by RNN. We trained and evaluated our model on the genome-wide PAS usage measurement obtained from 3’-mRNA sequencing of fibroblast cells from two mouse strains as well as their F1 hybrid. Our model, DeeReCT-APA, outperforms the state-of-the-art PAS quantification methods on the tasks that they are trained for, including pairwise comparison, highest usage prediction and ranking task. In addition, we demonstrated that modeling all the PAS of a gene simultaneously captures the mechanistic competition among the PAS and reveals the genetic variants with regulatory effects on PAS usage.

A similar idea of using BiLSTM to model competitive biological processes was proposed recently in [39]. The researchers used BiLSTM to model the usage level of competitive alternative 5’/3’ splice sites. Given the similarity of modeling competing polyadenylation sites and splice sites, it is therefore not surprising that DeeReCT-APA, which also incorporates BiLSTM to model the interactions among competing polyadenylation sites, achieves the best performance on the PAS quantification task.

Although DeeReCT-APA provides the first-of-its-kind method to model all the PAS of a gene, it still has room for improvement. As shown in Figure 3, the model has limited accuracy when the usage is very high or very low (comparing Figure 3B and Figure 3C). In addition, for allelic comparison as shown in Figure 5, some PAS with high allelic usage difference are predicted to be of low difference (false negatives, along X axis) and vice versa (false positives, along Y axis). One of the main reasons for our model’s limitation, as well as for all the other PAS quantification methods, is that all the existing genome-wide PAS quantification datasets used as training data could only sample the limited number of naturally occurring sequence variants. Although in our study the two parental strains from which the F1 hybrid mouse was derived are already the evolutionarily most distant ones among all the 17 mouse strains with complete genomic sequences, the number of genetic variants is still rather limited. To address this limitation and provide a complementary dataset, we are working on establishing a large-scale synthetic APA mini-gene reporter-based system which samples the regulatory effect of millions of random sequences (manuscript in preparation). Another limitation of our current model lies in the fact that it does not take all the factors with potential PAS regulatory effects into consideration. For example, transcription kinetics, i.e., the elongation rate of Pol II, which is not considered by the model in this study, can also affect APA choice [40]. Similarly, DeeReCT-APA does not take the distance between consecutive PAS into account, which, together with the transcription elongation rate, can also affect APA [41]. All of them are potential directions for further improvement.

Finally, very recently, Zhang et al. showed that effectively combining the power of deep learning and the information in RNA-seq data can significantly boost the performance for investigating the pattern of alternative splicing [42]. Indeed, our preliminary results showed that also for the recognition of APA patterns, there are substantial cases in which deep learning cannot make accurate prediction but utilizing the pattern of RNA-seq coverage around the cleavage site could provide clear evidence, and vice versa. Future work integrating the strength of deep learning on genomic sequences and experimental RNA-seq data will for certain not only improve the model performance, but also shed more light on the APA regulatory mechanisms.

## Data Availability

Our implementation of DeeReCT-APA using the PyTorch [37] library is available at the repository (https://github.com/lzx325/DeeReCT-APA-repo). The genome-wide PAS quantification dataset of parental and F1 mouse fibroblast cell is available in the subfolder ‘APA_ML’. As provided in [31], the raw sequencing data from which this dataset is derived is accessible at European Nucleotide Archive (http://www.ebi.ac.uk/ena) under the accession number PRJEB15336 (URL: https://www.ebi.ac.uk/ena/browser/view/PRJEB15336).

## Authors’ contributions

ZL, YH, WC, and XG conceived the project. ZL developed the deep learning model and did the computational experiments. Yisheng Li and BZ provided and pre-processed the dataset. JZ, XZ, and MZ provided additional biological insights on the experimental results. ZL, YH, WC, and XG drafted the paper. ZL, Yisheng Li, BZ, Yu Li, Yongkang Long, JZ, XZ, MZ, YH, WC, and XG read and approved the final manuscript.

## Acknowledgements

This work was supported by the King Abdullah University of Science and Technology (KAUST) Office of Sponsored Research (OSR) under Awards No. URF/1/4098-01-01, BAS/1/1624-01, FCC/1/1976-18-01, FCC/1/1976-23-01, FCC/1/1976-25-01, FCC/1/1976-26-01, and FCS/1/4102-02-01; by the International Cooperation Research Grant [No. GJHZ20170310161947503] from Science and Technology Innovation Commission of Shenzhen Municipal Government (YH); and by The Shenzhen Science and Technology Program (Grant No. : KQTD20180411143432337) (YH and WC).

## Supplementary material

### Supplementary File

DeeReCT-APA-Supplementary-File.pdf

**Figure S1 The structures of DeeReCT-APA models used in the ablation study**.

**A**. The structure of DeeReCT-APA with interaction layers but without BiLSTM. **B**. The structure of DeeReCT-APA with interaction layers removed. Comparing **A** with Figure 1 in the main text, it has BiLSTM removed and only has the affine layer in the interaction layers. In **B**, the interaction layers are removed altogether and DeeReCT-APA resorted to comparison-based training (to predict which one of the two PAS is of higher usage). Note that an additional affine layer is added on top of the Base Networks to cast the output of the base network (which is a vector) into a scalar.

**Figure S2 Comparison of the allelic usage difference prediction of DeeReCT-APA and Polyadenylation Code**.

F1 model fine-tuned from SP parental model is used. **A. B**. The horizontal axis is the ground truth allelic usage value difference between two homologous PAS (which is the BL usage value minus the SP usage value). The vertical axis shows the predicted allelic usage value difference. The scatter plot of DeeReCT-APA is shown in Panel **A** and Polyadenylation Code is shown in Panel **B**. As DeeReCT-APA predicts the usage value in percentage, we draw a red line that shows the perfect prediction. **C**. Pearson correlations between two quantities at different minimum allelic usage difference are shown in the table below.

**Figure S3 Visualization of convolutional filters in layer 1 of DeeReCT-APA**.

There are 40 convolutional filters in layer 1 of DeeReCT-APA. The model is trained on parental BL dataset and fine-tuned on F1.

**Figure S4 Visualization of convolutional filters in layer 2 of DeeReCT-APA**.

There are 40 convolutional filters in layer 2 of DeeReCT-APA. The model is trained on parental BL dataset and fine-tuned on F1.

**Table S1 List of features used in Feature-Net and their corresponding dimensions**.

**Table S2 List of hyperparameters for the three DeeReCT-APA models**.

**Table S3 Performance summary for the BL parental model and the F1 model fine-tuned from the BL parental model**.

**Table S4 Performance summary for the SP parental model and the F1 model fine-tuned from the SP parental model**.

**Table S5 Replicated Experiments of 5-fold cross validation on 5 random splits. Table S6 Comparison accuracy on dataset from [20]**

**Table S7 Replicated Experiments of ablation study**.

