## Supplementary Files for "DeeReCT-APA: Prediction of Alternative Polyadenylation Site Usage Through Deep Learning"

### Supplementary Materials for DeeReCT-APA: Prediction of Alternative Polyadenylation Site Usage Through Deep Learning

### S1 Feature extraction in Feature-Net

We in this section give a list of all the features extracted from raw sequence by the Feature-Net. A list of feature types and their dimensions are shown in Table S1. We also list out the polyadenylation signals, the AUE, CUE and ADE elements, as well as RBP motifs that are used:

Polyadenylation Signals: AATAAA, ATTAAA, TATAAA, AGTAAA, AAGAAA, AATATA, AATACA, CATAAA, GATAAA, AATGAA, TTTAAA, ACTAAA, AATAGA, AAAAAG, AAAATA, GGGGCT, AAAAAA, ATAAAA, AAATAA, ATAAAT, TTTTTT, ATAAAG, TAAAAA, CAATAA, TAATAA, ATAAAC

AUE Elements: GGGGAG, GUGGGG, GGGUGG, UUUGUA, GUAUUU, CUGUGU, UAUAUA, AUAUAU, UUUAUA, UGUAUA, AUGUAU, UGUAUU

CUE Elements: UAUUUU, UGUUUU, UUUUUU, AAUAAA, AUAAAG, AAAUAA

CDE Elements: GUGUCU, CUGCCU, UGUCUC, UUAUUU, UUUCUU, UGUUUU, UGUGUG, GUGUGU, CUGUGU, CUGGGG, UGUCUG, GUCUGU

ADE Elements: CCUCCC, CUCCCC, CACCCC, CCCGCC, CCCCGC, CCCGCG, GGUGGG, GGCUGG, GGGUGG, GGGCAG, GGCCAG, GGGGCC, GGGAGG, GGAGGG, GGGGAG

RBP Motifs: UUUUAU, GGGAGG, GGAGGG, GCUUGC, YGCY, YGCUKY, ARAAGA, UUUUCU, UCAY, CCWWHCC, CCYYCCH, UGGGRAD, GGGA, UKKGGK, GGSKG, UGUA, UGUGU, GAAGAA

### S2 The hyperparameters for DeeReCT-APA

We provide a full list of hyperparameters in Table S2. To limit the search space of hyperparameters, we choose to randomly sample hyperparameters that the model is most sensitive to, *i.e.* batch size, learning rate, L2 weight decay and dropout rate and choose the combination that makes the model perform best on the validation set. Note that since we use three different designs for Base-Net, their best hyperparameters are slightly different. Note the best values for hyperparameters that affect regularization of model, *i.e.* L2 weight decay and dropout rate, are higher for more complex model (Multi-Conv-Net), which is reasonable, as complex models are more prone to overfitting and thus their capacity should be limited more by regularization.

### S3 Performance of SP parental model and F1 model fine-tuned from SP parental model

As in the main text, we provide the overall performance measures for SP parental models and F1 models fine-tuned from SP parental models for the three different models: DeeReCT-APA, Polyadenylation Code and DeepPASTA (Table S4). The DeeReCT-APA with Multi-Conv-Net shows generally better performance than Polyadenylation Code and DeepPASTA.

We also globally show the quality of prediction of DeeReCT-APA and Polyadenylation Code in F1 hybrid cell using the scatter plot of predicted allelic usage difference versus ground truth allelic usage difference, as in the main text. As it is shown in Figure S2, DeeReCT-APA shows higher correlation between predicted and ground truth allelic usage difference than Polyadenylation Code.

### S4 Additional performance measures

In this section, we provide results of some additional tests on DeeReCT-APA's performance. The first is to test the significance of DeeReCT-APA's improvement over existing methods. The second is to use a benchmark dataset other than the one from [1] to evaluate DeeReCT-APA's comparison accuracy.

To test the significance of DeeReCT-APA's improvment over Polyadenylation Code and DeepPASTA, we conducted 5 replicated experiments, where in each of them we independently split the dataset into 5 folds and perform 5-fold cross validation on them. With completely identical settings, we train the model on parental dataset and fine-tune on F1. We report the comparison accuracy of each method in different replicates seperately (Table S7). The t-tests between DeeReCT-APA and Polyadenylation Code as well as DeeReCT-APA and DeepPASTA show that the improvement of DeeReCT-APA over Polyadenylation Code and DeepPASTA is significant (last column).

We also show the performance of DeeReCT-APA on the dataset obtained from [2]. Since this is a multi-tissue dataset, we report the comparison accuracy in each tissue separately. One can see that DeeReCT-APA outperforms existing methods in most of them.

### Supplementary Figures and Tables

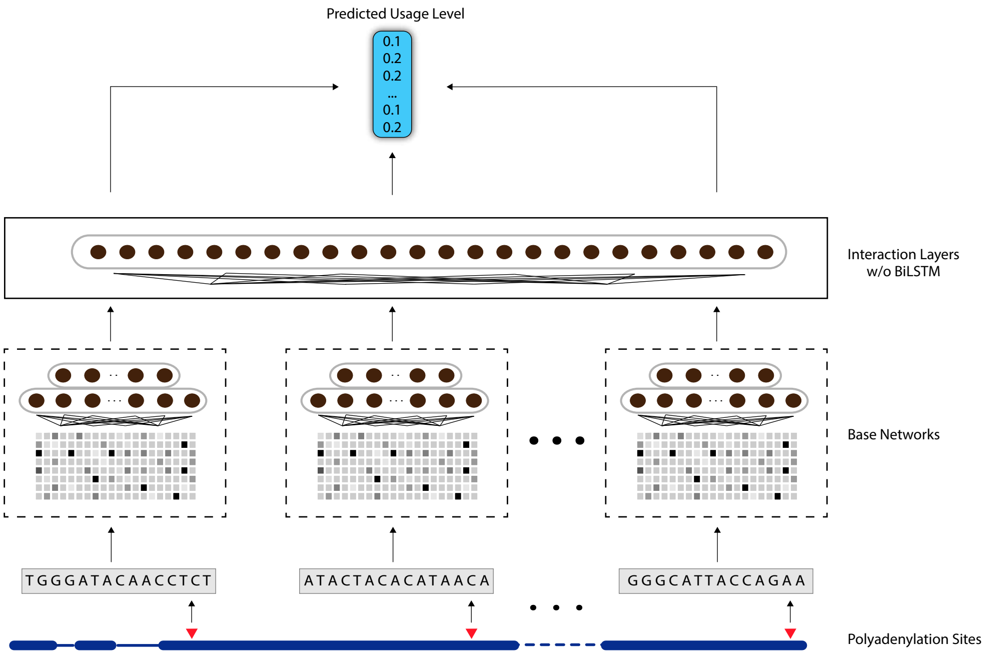


(a) The structure of DeeReCT-APA with interaction layers but without BiLSTM

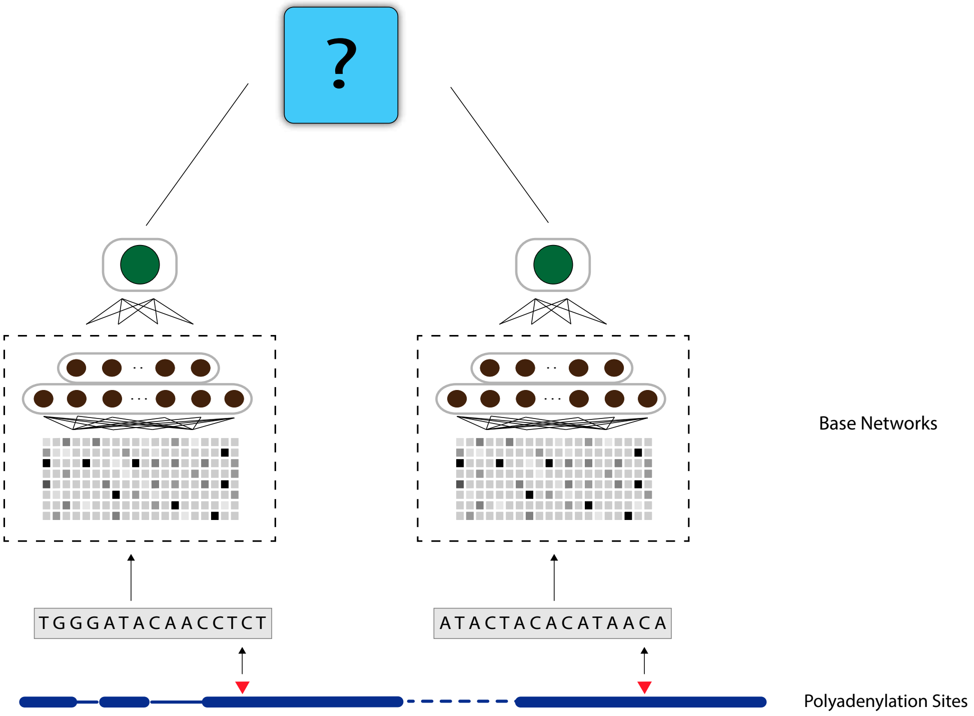


(b) The structure of DeeReCT-APA with interaction layers removed

Figure S1: The structures of DeeReCT-APA models used in the ablation study.
Comparing (a) with Figure 1 in the main text, it has BiLSTM removed and only has the affine layer in the interaction layers. In (b), the interaction layers are removed altogether and DeeReCT-APA resorted to comparison-based training (to predict which one of the two PAS is of higher usage). Note that an additional affine layer is added on top of the Base Networks to cast the output of the base network (which is a vector) into a scalar.

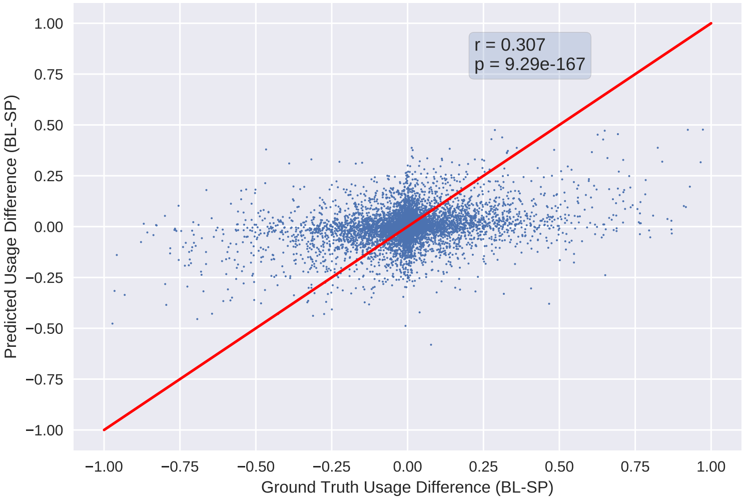


(a) DeeReCT-APA

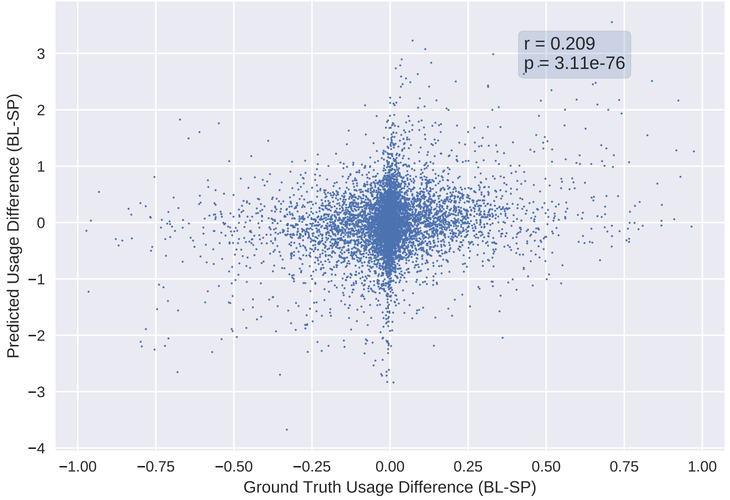


(b) Polyadenylation Code

| Min. Allelic Usage Difference | | 0.1 | 0.2 | 0.3 | 0.4 | 0.6 | 0.8 |
|---|---|---|---|---|---|---|---|
| DeeReCT-APA | Pearson R | 0.400 | 0.467 | 0.507 | 0.536 | 0.645 | 0.708 |
| | p value | $3.5 \times 10^{-103}$ | $3.5 \times 10^{-74}$ | $4.9 \times 10^{-48}$ | $5.2 \times 10^{-31}$ | $1.5 \times 10^{-16}$ | $4.7 \times 10^{-5}$ |
| Polyadenylation Code | Pearson R | 0.307 | 0.351 | 0.366 | 0.402 | 0.465 | 0.568 |
| | p value | $4.4 \times 10^{-57}$ | $5.3 \times 10^{-39}$ | $2.3 \times 10^{-27}$ | $5.0 \times 10^{-19}$ | $3.2 \times 10^{-9}$ | $7.2 \times 10^{-3}$ |

(c) Pearson Correlation of the prediction of DeeReCT-APA and Polyadenylation Code at different minimum allelic usage difference

Figure S2: Comparison of the allelic usage difference prediction of DeeReCT-APA and Polyadenylation Code.
F1 model fine-tuned from SP parental model is used. The horizontal axis is the ground truth allelic usage value difference between two homologous PAS (which is the BL usage value minus the SP usage value). The vertical axis shows the predicted allelic usage value difference. As DeeReCT-APA predicts the usage value in percentage, we draw a red line that shows the perfect prediction. Pearson correlations between two quantities at different minimum allelic usage difference are shown in the table below.

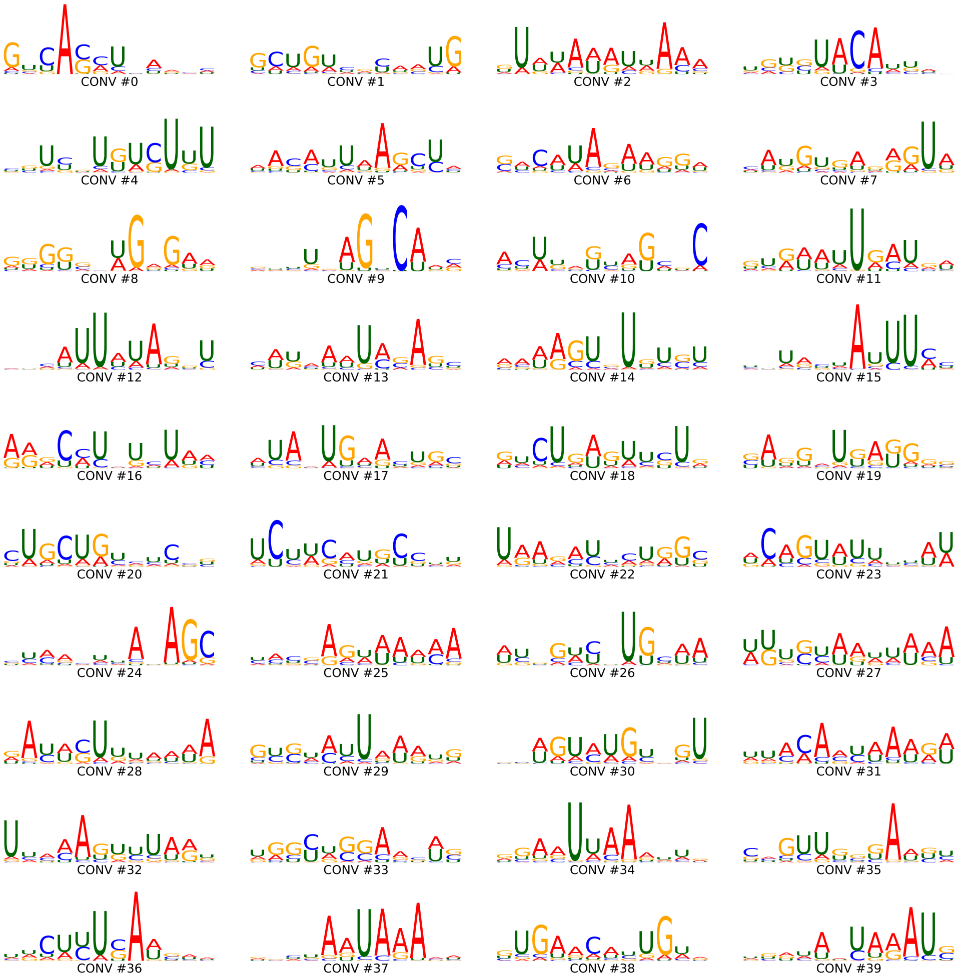


Figure S3: Visualization of convolutional filters in layer 1 of DeeReCT-APA.
There are 40 convolutional filters in layer 1 of DeeReCT-APA. The model is trained on parental BL dataset and fine-tuned on F1.

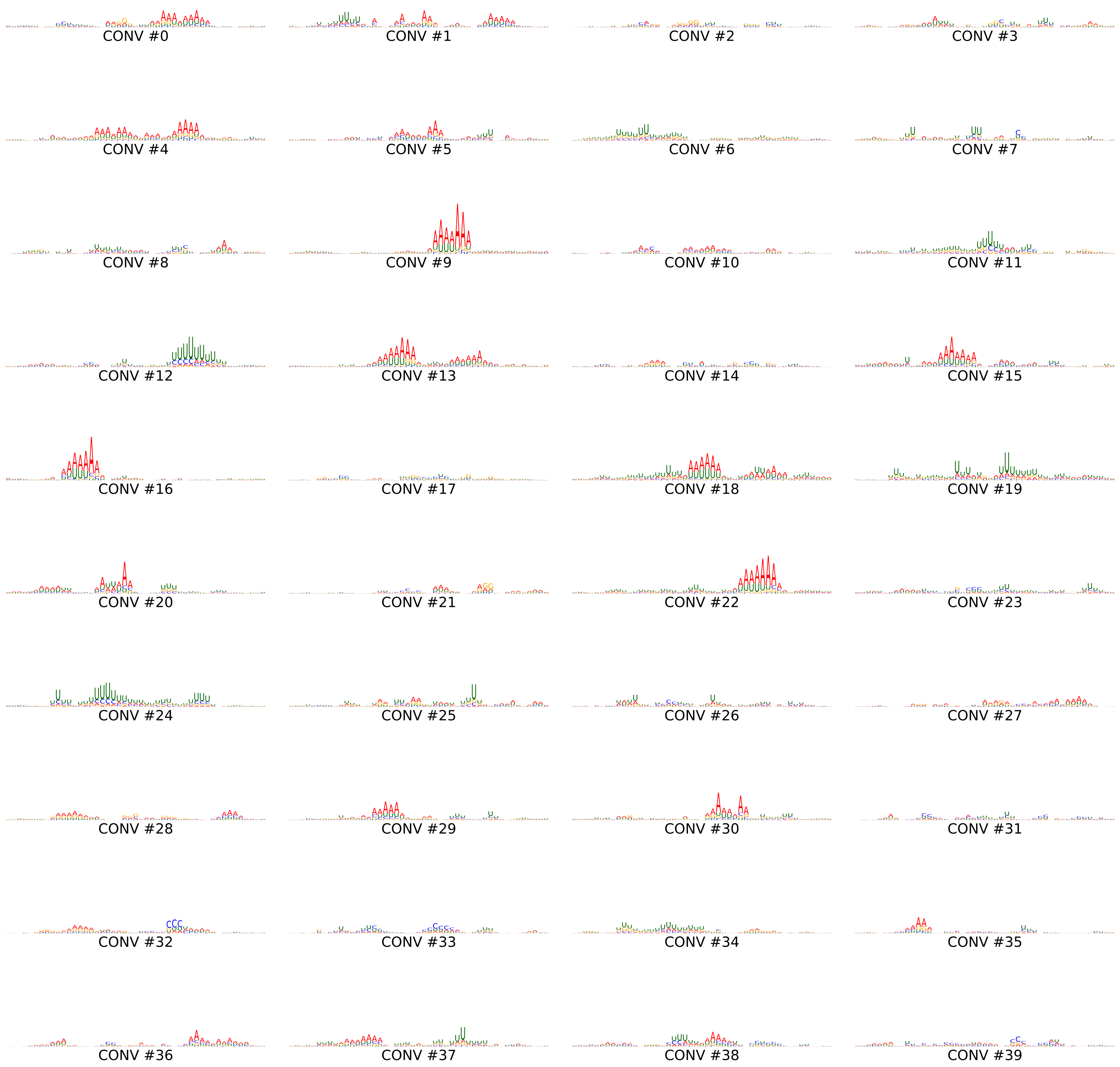


Figure S4: Visualization of convolutional filters in layer 2 of DeeReCT-APA.
There are 40 convolutional filters in layer 2 of DeeReCT-APA. The model is trained on parental BL dataset and fine-tuned on F1.

| Feature | Number of dims |
|---|---|
| Polyadenylation Signals | 52 |
| AUE Elements | 12 |
| CUE Elements | 2 |
| ADE Elements | 12 |
| RBP Motifs | 72 |
| 1-mers | 16 |
| 2-mers | 64 |
| 3-mers | 256 |
| 4-mers | 992 |
| Nucleosome Occupancy | 12 |
| Position | 1 |

Table S1: List of features used in Feature-Net and their corresponding dimensions.

| Hyperparameter | DeeReCT-APA (Single-Conv-Net) | DeeReCT-APA (Multi-Conv-Net) | DeeReCT-APA (Feature-Net) |
|---|---|---|---|
| Batch size {16,32,64} | 32 | 32 | 32 |
| Learning rate [1e-5,1e-2] | 1e-3 | 1e-2 | 2e-5 |
| L2 weight decay [1e-5,1e-2] | 2e-5 | 1e-3 | 1e-5 |
| Conv filter width (layer 1) | 12 | 12 | - |
| Number of conv filters (Layer 1) | 16 | 40 | - |
| Pool width (Layer 1) | 12 | 3 | - |
| Conv filter width (Layer 2) | - | 12 | - |
| Number of conv filters (Layer 2) | - | 40 | - |
| Pool width (Layer 2) | - | 4 | - |
| LSTM input dim | 200 | 200 | 200 |
| LSTM output dim | 100 | 100 | 100 |
| Dropout rate [0.1,0.9] | 0.2 | 0.7 | 0.2 |

Table S2: List of hyperparameters for the three DeeReCT-APA models.
For the hyperparameters that are sampled randomly, the ranges (in brackets) or sets (in braces) that they are sampled from are shown. The hyperparameters that are not applicable for a specific model are denoted by "-".

| Model | Performance on Parental Dataset | | | |
|---|---|---|---|---|
| | MAE* | Comparison Accuracy* | Highest Usage Prediction Accuracy* | Averaged Spearman's Correlation |
| DeeReCT-APA (Feature-Net) | 17.60% ± 0.3% | 76.90% ± 0.9% | 61.50% ± 1.3% | 0.4999 ± 0.014 |
| DeeReCT-APA (Single-Conv-Net) | 17.42% ± 0.3% | 77.20% ± 0.7% | 62.22% ± 0.5% | 0.5035 ± 0.011 |
| DeeReCT-APA (Multi-Conv-Net) | **17.22% ± 0.3%** | **77.64% ± 0.4%** | **63.48% ± 0.9%** | **0.5140 ± 0.021** |

* The values for a random predictor are 43.12%, 50.00% and 25.49% respectively.

(a) Performance on the Parental Dataset (BL)

| Model | Performance on F1 Dataset | | | |
|---|---|---|---|---|
| | MAE* | Comparison Accuracy* | Highest Usage Prediction Accuracy* | Averaged Spearman's Correlation |
| DeeReCT-APA (Feature-Net) | 18.20% ± 0.6% | 76.30% ± 1.6% | 61.10% ± 1.9% | 0.4661 ± 0.025 |
| DeeReCT-APA (Single-Conv-Net) | 18.60% ± 0.6% | 74.94% ± 2.4% | 61.68% ± 1.8% | 0.4602 ± 0.027 |
| DeeReCT-APA (Multi-Conv-Net) | **17.80% ± 0.4%** | **77.14% ± 1.2%** | **64.52% ± 0.7%** | **0.4957 ± 0.009** |

* The values for a random predictor are 40.96%, 50.00% and 28.56% respectively.

(b) Performance on the F1 Dataset (fine-tuned from parental BL model)

Table S3: Performance summary for the BL parental model and the F1 model fine-tuned from the BL parental model. The table shows the performance of DeeReCT-APA with three different Base-Nets across four evaluation metrics. Results are shown in $mean \pm std$ format.

| Model | Performance on Parental Dataset | | | |
|---|---|---|---|---|
| | MAE* | Comparison Accuracy* | Highest Usage Prediction Accuracy* | Averaged Spearman's Correlation |
| DeeReCT-APA (Multi-Conv-Net) | **17.90% ± 0.2%** | **76.20% ± 1.0%** | **61.58% ± 1.4%** | **0.4930 ± 0.018** |
| Polyadenylation Code | N/A | 74.98% ± 2.1% | 59.60% ± 2.4% | 0.4610 ± 0.026 |
| DeepPASTA | N/A | 72.12% ± 0.9% | 56.98% ± 1.3% | 0.4273 ± 0.011 |

* The values for a random predictor are 43.27%, 50.00% and 25.49% respectively.

(a) Performance on Parental Dataset (SP)

| Model | Performance on F1 Dataset | | | |
|---|---|---|---|---|
| | MAE* | Comparison Accuracy* | Highest Usage Prediction Accuracy* | Averaged Spearman's Correlation |
| DeeReCT-APA (Multi-Conv-Net) | **17.72% ± 0.4%** | **76.52% ± 1.5%** | **63.96% ± 1.1%** | **0.4904 ± 0.007** |
| Polyadenylation Code | N/A | 74.62% ± 2.5% | 58.62% ± 1.8% | 0.4246 ± 0.030 |
| DeepPASTA | N/A | 69.22% ± 0.79% | 53.18% ± 2.7% | 0.3537 ± 0.029 |

* The values for a random predictor are 40.96%, 50.00% and 28.56% respectively.

(b) Performance on F1 Dataset (fine-tuned from parental SP model)

Table S4: Performance summary for the SP parental model and the F1 model fine-tuned from the SP parental model. The table shows the performance of the three models across four evaluation metrics. Results are shown in $mean \pm std$ format.

| Model | Comparison Accuracy on Parental Dataset | | | | | |
|---|---|---|---|---|---|---|
| | Replicate 1 | Replicate 2 | Replicate 3 | Replicate 4 | Replicate 5 | p-value |
| DeeReCT-APA (Multi-Conv-Net) | 77.64% | 77.53% | 77.49% | 77.92% | 77.34% | - |
| Polyadenylation Code | 75.88% | 75.39% | 76.01% | 75.77% | 75.69% | $1.6 \times 10^{-4}$ |
| DeepPASTA | 74.08% | 73.94% | 74.12% | 74.20% | 73.87% | $4.5 \times 10^{-7}$ |

(a) Replicated Experiments on Parental Dataset (BL)

| Model | Comparison Accuracy on F1 Dataset | | | | | |
|---|---|---|---|---|---|---|
| | Replicate 1 | Replicate 2 | Replicate 3 | Replicate 4 | Replicate 5 | p-value |
| DeeReCT-APA (Multi-Conv-Net) | 77.14% | 77.94% | 76.88% | 77.09% | 77.10% | - |
| Polyadenylation Code | 74.20% | 74.23% | 74.15% | 74.02% | 74.31% | $6.5 \times 10^{-5}$ |
| DeepPASTA | 70.14% | 70.20% | 70.87% | 71.02% | 70.46% | $3.1 \times 10^{-5}$ |

(b) Replicated Experiments on F1 Dataset

Table S5: Replicated Experiments of 5-fold cross validation on 5 random splits.
The table shows the averaged comparison accuracy across the 5-fold cross validation of the three models on parental BL dataset and F1 dataset. There are 5 replicates for the experiment. At the end of each row, the table shows the p-value of the t-test of DeeReCT-APA's performance compared against the model of that row.

| Tissue | Comparison Accuracy | | |
|---|---|---|---|
| | DeepPASTA | Polyadenylation Code | DeeReCT-APA (Multi-Conv-Net) |
| Brain | **0.908** | 0.895 | 0.895 |
| Breast | **0.900** | 0.886 | **0.900** |
| ES Cell | 0.910 | 0.911 | **0.925** |
| Ovary | 0.903 | 0.895 | **0.912** |
| SK Muscle | 0.906 | 0.893 | **0.914** |
| Testis | 0.893 | 0.856 | **0.905** |
| BCells1 | **0.905** | 0.896 | 0.904 |
| BCells2 | 0.901 | 0.893 | **0.907** |

Table S6: Comparison accuracy on dataset from [2].
The performance of Polyadenylation Code and DeepPASTA is obtained from [2] and [3].

| Model | Comparison Accuracy on Parental Dataset | | | | | |
|---|---|---|---|---|---|---|
| | Replicate 1 | Replicate 2 | Replicate 3 | Replicate 4 | Replicate 5 | p-value |
| DeeReCT-APA(Multi-Conv-Net) (No Interaction Layer) | 76.12% | 76.04% | 76.27% | 76.13% | 76.29% | - |
| DeeReCT-APA(Multi-Conv-Net) (w/o BiLSTM) | 77.12% | 77.00% | 77.21% | 77.29% | 76.94% | $2.5 \times 10^{-6}$ |
| DeeReCT-APA(Multi-Conv-Net) (BiLSTM) | 77.64% | 77.53% | 77.49% | 77.92% | 77.34% | $3.7 \times 10^{-3}$ |

(a) Replicated Experiments on Parental Dataset (BL)

| Model | Comparison Accuracy on F1 Dataset | | | | | |
|---|---|---|---|---|---|---|
| | Replicate 1 | Replicate 2 | Replicate 3 | Replicate 4 | Replicate 5 | p-value |
| DeeReCT-APA(Multi-Conv-Net) (No Interaction Layer) | 76.28% | 76.33% | 76.35% | 76.21% | 76.15% | - |
| DeeReCT-APA(Multi-Conv-Net) (w/o BiLSTM) | 76.77% | 76.88% | 76.87% | 77.16% | 76.49% | $1.1 \times 10^{-3}$ |
| DeeReCT-APA(Multi-Conv-Net) (BiLSTM) | 77.14% | 77.94% | 76.88% | 77.09% | 77.10% | $9.9 \times 10^{-2}$ |

(b) Replicated Experiments on F1 Dataset

Table S7: Replicated Experiments of ablation study.
The table shows averaged comparison accuracy across 5-fold cross validation of the two ablated models and the full DeeReCT-APA model on parental BL dataset and F1 dataset. There are 5 replicates for the experiment. At the end of each row, the table shows p-value of t-test of *the current row's* performance compared against the model of *the previous row*. In terms of most replicates, there is improvement over the previous row.

[3] Ashraful Arefeen, Tao Jiang, and Xinshu Xiao. Deeppasta: Deep neural network based polyadenylation site analysis. 2019.
